# Children with cystic fibrosis are infected with multiple subpopulations of *Mycobacterium abscessus* with different antimicrobial resistance profiles

**DOI:** 10.1101/436261

**Authors:** Liam P. Shaw, Ronan M. Doyle, Ema Kavaliunaite, Helen Spencer, Francois Balloux, Garth Dixon, Kathryn A. Harris

## Abstract

**Background:** Children with cystic fibrosis (CF) can develop life-threatening infections of *Mycobacterium abscessus*. These present a significant clinical challenge, particularly when the strains involved are resistant to antibiotics. Recent evidence of within-patient subclones of *M. abscessus* in adults with CF suggests the possibility that within-patient diversity may be relevant for the treatment of pediatric CF patients.

**Methods:** We performed whole genome sequencing (WGS) on 32 isolates of *M. abscessus* from multiple body sites for two patients with CF undergoing treatment at Great Ormond Street Hospital, UK, in 2015.

**Results:** We found evidence of extensive diversity within patients over time. Clustering analysis of single nucleotide variants (SNVs) revealed that each patient harboured multiple subpopulations, which were differentially abundant between sputum, lung samples, chest wounds, and pleural fluid. Sputum isolates did not reflect overall within-patient diversity, including failing to detect subclones with mutations previously associated with macrolide resistance (*rrl* 2058/2059). Some variants were present at intermediate frequencies before lung transplant. The time of transplant coincided with extensive variation, suggesting that this event is particularly disruptive for the microbial community, but transplant did not clear the *M. abscessus* infection and both patients died as a result of this infection.

**Conclusions:** Isolates of *M. abscessus* from sputum do not always reflect the entire diversity present within the patient, which can include subclones with differing AMR profiles. Awareness of this phenotypic variability, with sampling of multiple body sites in conjunction with WGS, may be necessary to ensure the best treatment for this vulnerable patient group.

**Key point summary:**

- Children with cystic fibrosis undergoing lung transplant harbour multiple subpopulations of *M. abscessus*.
- Subpopulations can have different antimicrobial resistance genotypes.
- Sputum isolates do not reflect the genetic diversity within a patient.

## Introduction

*Mycobacterium abscessus* is a nontuberculous mycobacteria (NTM) which has recently emerged as a major pathogen in cystic fibrosis (CF) patients^1^. Infection with *M. abscessus* is associated with poor clinical outcomes, particularly in conjunction with lung transplantation^2^. Treatment is challenging due to the intrinsic resistance of *M. abscessus* to many classes of antibiotics^3^, along with certain genotypes drastically altering the efficacy of antibiotic courses^4^. The antimicrobial resistance (AMR) profile of isolates is highly relevant for treatment, but current diagnostic work mainly uses isolates from sputum, which may not reflect the full range of genetic diversity within the patient and therefore fail to recover the true AMR profile.

Minority variants from WGS have been used to infer the presence of multiple subpopulations (subclones) in longitudinal sputum isolates of *M. abscessus*^5^. However, it remains an open question whether patients harbour further genetic diversity within the lung that is not sampled by sputum. The lung is known to be capable of harbouring considerable pathogen diversity in chronic infections. For example, *M. tuberculosis* has been shown to exist as multiple subpopulations with different AMR profiles^6^. Furthermore, the relevance of this genetic diversity for treatment is not yet known.

For this reason, we investigated longitudinal isolates from two patients infected with *M. abscessus subsp. abscessus* undergoing lung transplant at Great Ormond Street Hospital (Figure 1), and identified variable genomic positions within samples (single nucleotide variants, SNVs). By including isolates from sputum samples but also biologically important compartments such as pleural fluid, lung tissue, and swabs from chest wounds, we aimed to establish the extent and significance of within-patient variation in *M. abscessus* for this vulnerable group.

**Figure 1.**
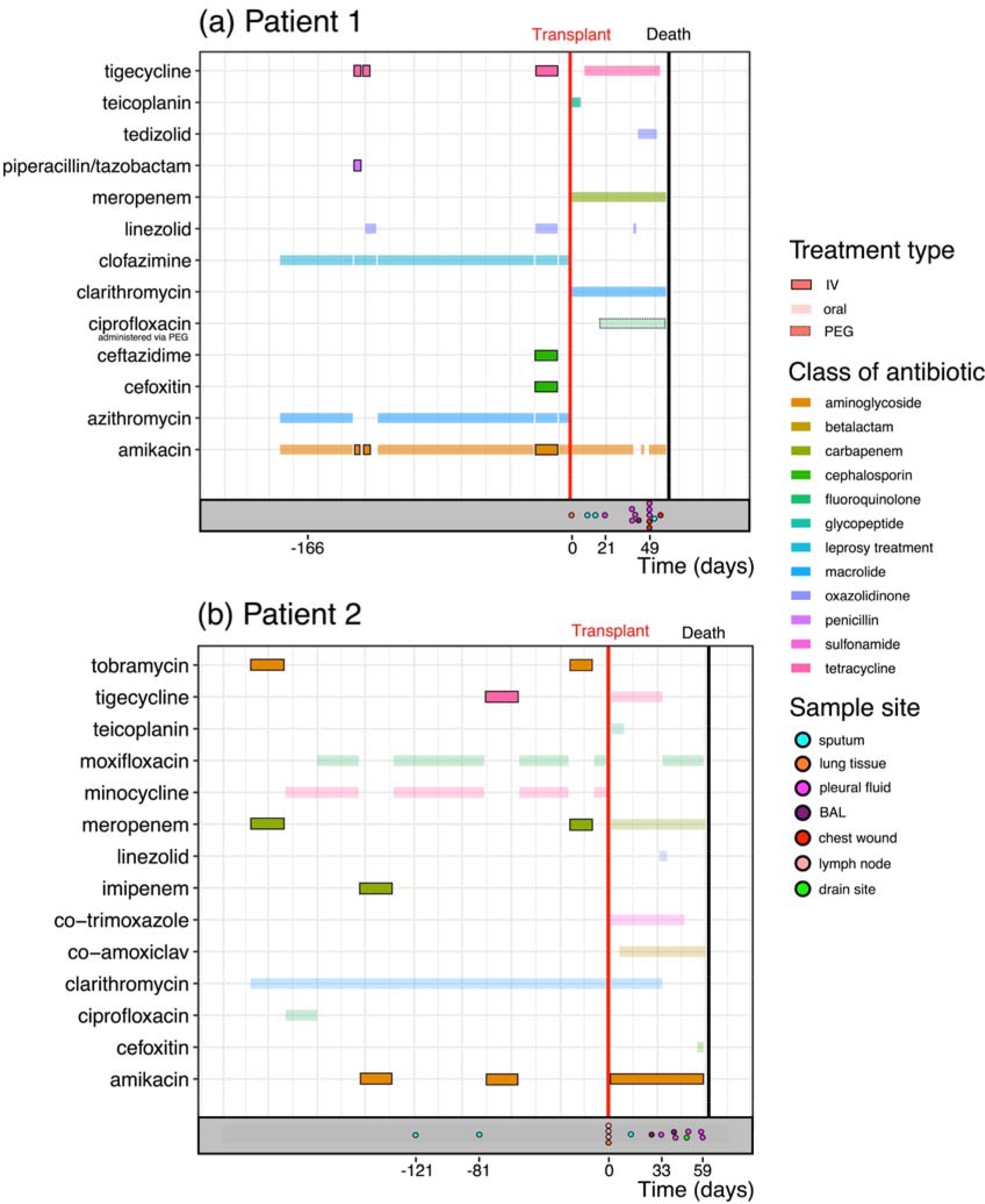
Overview of the sampling scheme and antibiotic treatment. Treatment regime including intravenous antibiotics (boxed coloured lines) and oral antibiotics (faint coloured lines) for both patients during the six-month period prior to lung transplant (red vertical line). The sampling scheme is represented at the bottom of each panel (coloured points on grey background).

## Methods

### Patient cohort and sample collection

Great Ormond Street Hospital (GOSH) is a large regional centre for paediatric CF patients and the largest paediatric lung transplant centre in the UK. The two patients in this study (Patient 1 and Patient 2) were from other CF centres and were seen at GOSH for a transplant assessment, during the lung transplant procedure and post-transplant (Table 1). Both patients in this study underwent regular respiratory microbiological diagnostic investigations, including specific stain and culture for mycobacteria on sputum pre- and post-transplant and explanted lung tissue, bronchoalveolar lavage, pleural fluid and clamshell incision wound swabs post-transplant. All further microbiological investigations were carried out on ‘sweeps’ from pure culture plates. All *M. abscessus* isolates cultured in our laboratory are identified to sub-species level by PCR and sequencing of *hsp65* and *rpoB* genes and inducible macrolide resistance predicted by PCR and sequencing of the *erm(41)* gene as previously described^7^. VNTR profiles were obtained for selected isolates as previously described^8^. Phenotypic sensitivity data was obtained from the mycobacterial reference laboratory (Table 3).

**Table 1.** Summary statistics for the de novo reference genomes of the two patients in this study. References were assembled for the first temporal sample from each patient (patient_1_S1 and patient_2_S1; see Methods).

|  | Patient 1 | Patient 2 |
| --- | --- | --- |
| Total bases | 5,172,759 | 5,275,491 |
| Mean depth of coverage | 25.3X | 74.1X |
| Number of contigs | 28 | 46 |

**Table 2.** patient clinical data. Dates are relative to the day of transplant.

| Patient | *M. abscessus* sub-species | Sex | CF genotype | Date of last smear-positive sample | Date of last *M. abscessus* isolate |
| --- | --- | --- | --- | --- | --- |
| Patient 1 | ABS | Female | F508del/F508del | -166 days | +56 days |
| Patient 2 | ABS | Male | F508del/W1282X | -81 days | +59 days |

**Table 3.**
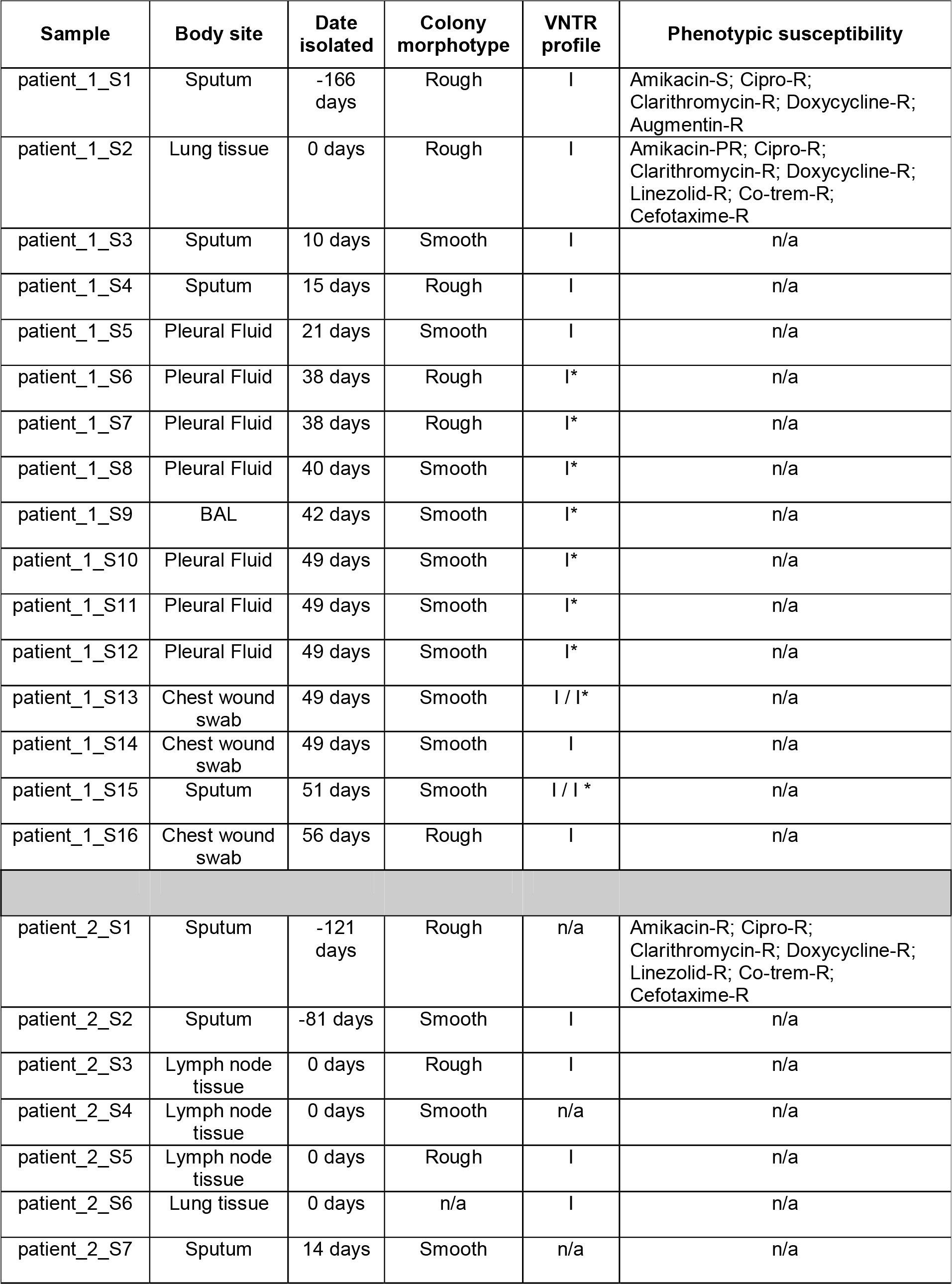
Routine microbiology data about isolates in this study. Dates are relative to the date of transplant for each patient. An asterisk (*) indicates that the VNTR profile differed at one locus. n/a indicates data not available. Phenotypic susceptibility was available for a minority of isolates (S=susceptible, R=resistant, PR=partially resistant).

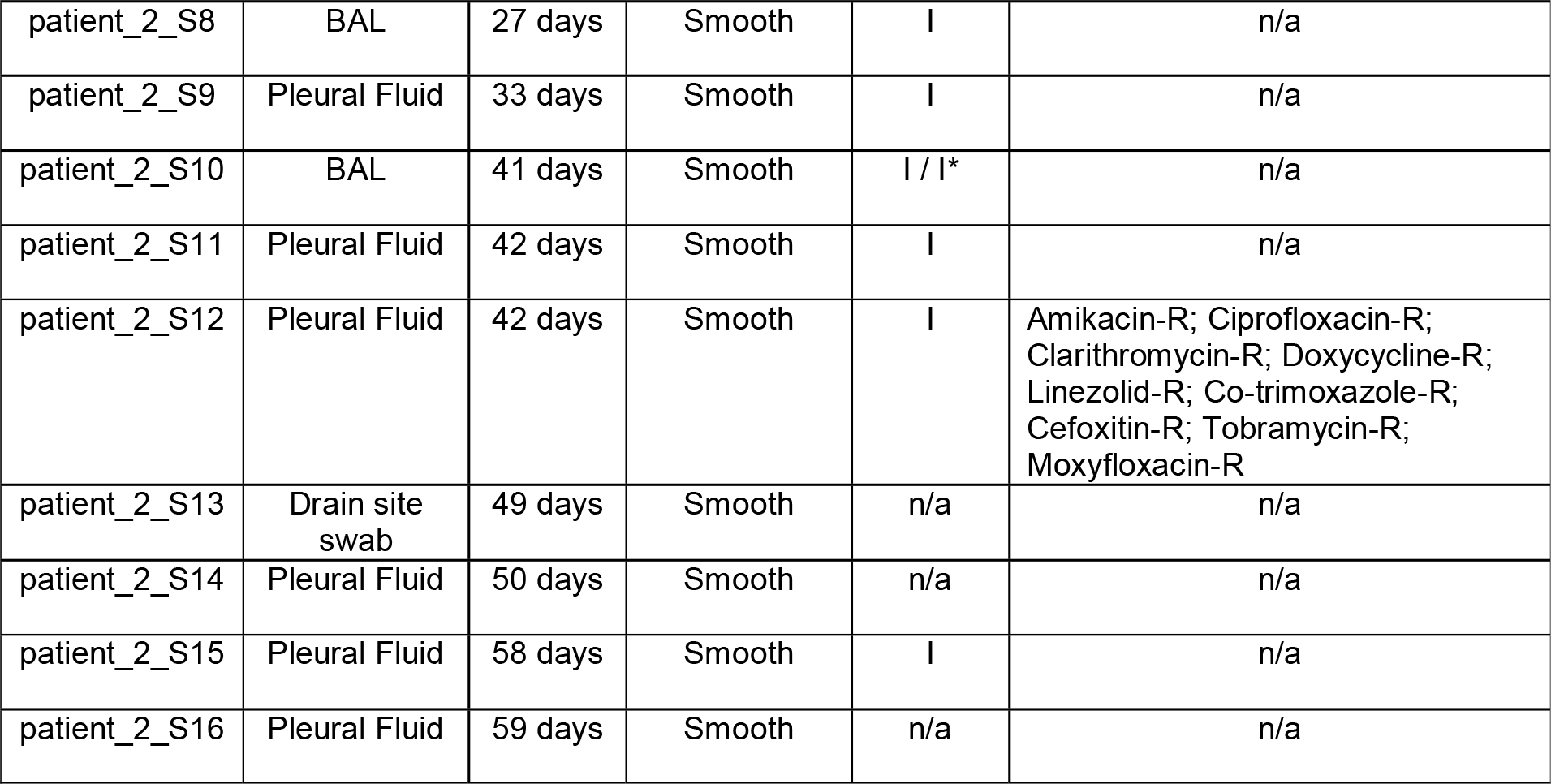

Demographic and clinical data were extracted from the Patient Administration System (PIMS) and microbiological data from the Laboratory Information Management system (OMNI-client ISS) using SQL (structured query language) databases and Excel spreadsheets. Additional sources of information included CF and transplantation databases. Details of antimicrobial therapy administered prior to transplantation was provided by the referring CF centres. All investigations were performed in accordance with the Hospitals Research governance policies and procedures.

### Whole genome sequencing

16 isolates from Patient 1 and 16 isolates from Patient 2 underwent DNA extraction as previously described^8^ with the addition of a bead beating step. Total DNA concentration was determined using the Qubit high-sensitivity (HS) assay kit (Thermofisher) and a sequencing library was prepared from 50 ng of DNA using the Nextera Library Preparation kit (Illumina). Post-PCR clean-up was carried out using Ampure XP beads (Beckman). Library size was validated using the Agilent 2200 TapeStation with Agilent D1000 ScreenTape System and 150bp paired-end reads were sequenced on a NextSeq 550 system (Illumina). Raw sequencing reads have been deposited on ENA (Study Accession PRJEB28875) as well as two assemblies used as ‘d*e novo* references’ (see below).

### Sequence data analysis

Initial variable nucleotide tandem repeat (VNTR) typing carried out as described previously^8,9^ suggested the possibility of mixed infections, based on results intermediate between VNTR I and the closely related VNTR I* profile (differing at one locus). A preliminary mapping of all isolates to the standard *M. abscessus* strain ATCC 19977 chromosome (NCBI Accession: CU458896.1) showed that the mean coverage at 10X was ~91%, contrasted with >99% for a representative set of VNTR II isolates from another patient sequenced according to the same protocol (not shown). In order to ensure we captured as much of the genetic diversity as possible, we therefore adopted a hybrid *de novo* and mapping approach. We selected the first isolate (temporally) for each patient and performed *de novo* assembly with Spades v3.10.0 with the --careful switch and otherwise default parameters^10^. After removing contigs with <10,000 bases to reduce the possibility of including small mobile genetic elements, this first *de novo* assembly was used as a new reference to map raw reads from other isolates using bwa mem v0.7.12 with default parameters^11^. This produced ‘d*e novo* references’ for Patient 1 and Patient 2 containing 5.17Mb and 5.28Mb respectively (Table 1). Contigs were reordered against the *M. abscessus* ATCC 19977 chromosome using Mauve v2.4.0 (2015-02-25)^12^.

### Variant identification and clustering analysis

In brief, the mapping file was sorted and indexed using picard v1.130, then GATK v3.30 was used to create a combined variant call format (VCF) file for each patient. Each position required a mapping depth > 30 in all samples from a patient to be included in downstream analysis. We manually inspected the ‘self-mapping’ of the reads from the first temporal sample to its own *de novo* assembly using IGV v2.4.10 ^13^ to identify small regions where mapping was problematic. We removed SNVs within isolated regions where the self-mapping had unexpected peaks in coverage (Patient 1: contigs 12 (12,575-13,519bp) and 19 (96,916-97,032bp); Patient 2: contig 17 (12,718-12,770bp) as well as SNVs where the reference allele fraction from the self-mapping reads was < 5%.

As noted by Bryant *et al.*^5^, patterns of linkage of variants in for *M. abscessus* can be suggestive of the existence of subpopulations. We aimed to establish a conservative lower bound for the number of clonal subpopulations within a patient, inferring their existence from the linkage patterns of variant frequencies across all samples. We note that patterns of linkage disequilibrium can also occur due to recombination. We therefore attempted to remove local recombination in our analysis. Using the SNVs obtained via the mapping and filtering methods described above, we hierarchically clustered SNVs using Ward’s minimum variance criterion^14^ applied to Euclidean distances between allele frequencies with a dissimilarity threshold of 1 to define clusters. We removed clusters containing <4 SNVs. We also removed putative local recombination regions by removing clusters where the SNVs were distributed within a total range < 100,000bp (corresponding to a threshold of ~2% of the total length of the *M. abscessus* genome).

In general, the inference of haplotype frequencies using variant frequencies from short sequencing reads for a microbial population undergoing recombination is a complex problem^15^. However, as we are not attempting to comment on abundances of subpopulations but only their presence, we did not need to infer haplotype frequencies. Observing *n* distinct clusters of variants within a patient over time (i.e. allele frequencies that co-vary in step with each other) means that there must be at least *n* bacterial haplotypes within the population producing these patterns. This fact holds even when recombination is present. Therefore, observing distinct clusters of linked variants tells us that distinct subpopulations of *M. abscessus* exist within individual samples.

## Results

### Individuals harbour extensive variation

We observed multiple positions in the *M. abscessus* genome which varied between different isolates within a patient over time (total variable positions used for clustering across all isolates, patient 1: 54 positions, patient 2: 64 positions), although isolates remained highly similar on average and were clearly the same infecting strain (mean inter-isolate SNV distances patient 1: 2.07 +/− 0.92 SNVs, patient 2: 1.96 +/− 1.81 SNVs). Subsets of these SNVs showed patterns of linkage across the *M. abscessus* genome (Figure 2a). Similar clustered patterns of linked SNVs could also arise due to recombination, but the final groups of clustered SNVs were spread across the genome suggesting that multiple subclones were present. Even if recombination were present, leading to mixtures of clusters (i.e. different haplotypes), then the observed abundance patterns still required multiple subclones. A neighbour-joining tree produced from distances between isolates based on these allele frequencies suggested the presence of multiple subclones (Figure 2b).

**Figure 2.**
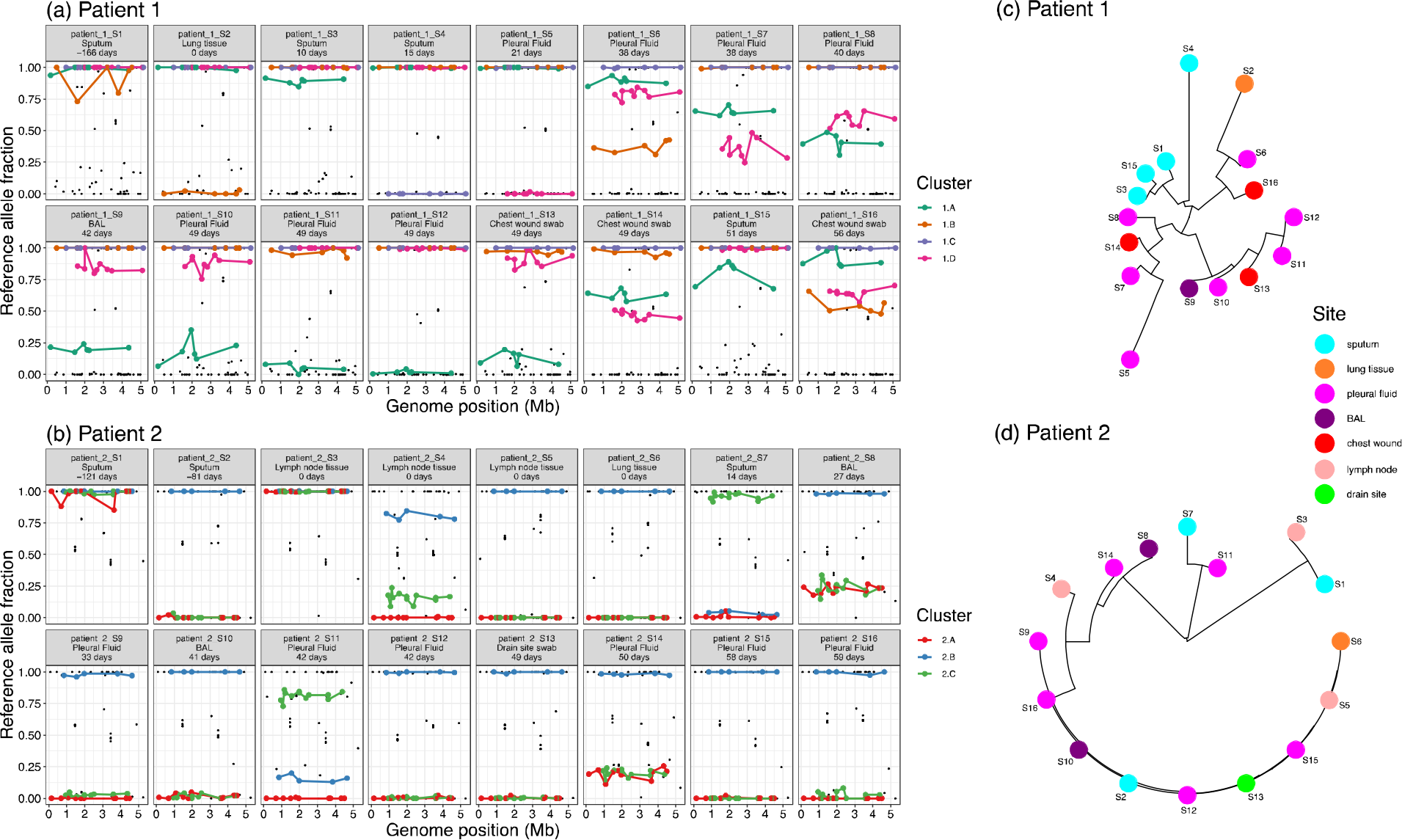
Linkage patterns of SNVs across samples suggest the presence of closely-related subpopulations within patients. **(a, b)** SNVs were grouped into clusters (colours) using an unsupervised clustering technique, showing clear patterns of abundance across samples (see Methods). Genome position was inferred by ordering *de novo* contigs against the *M. abscessus* ATCC 19977 reference genome. **(c, d)** Midpoint-rooted neighbour-joining tree based on Euclidean distances between samples using these clustered SNVs show this variation within patients over time (numbers) and body site (colours).

### Within-patient variation includes antimicrobial resistance mutations

Macrolide resistance in *M. abscessus* is driven by mutations at well-established positions in the *rrl* gene. Both patients received macrolides almost continuously throughout the six months prior to transplant (patient 1: oral azithromycin, patient 2: oral clarithromycin). While initial isolates taken earlier in treatment were susceptible, we observed that resistance alleles at these positions (C/G) increased in abundance over time (Figure 3). Notably, all sputum isolates from patient 1 showed a susceptible allele at position 2059 whereas isolates from pleural fluid and clamshell incision wound swabs carried a resistance allele (A2059C, Figure 3a). There was also substantial variation within sets of isolates taken on the same day. For example, three isolates from different samples of lymph node tissue taken on the day of transplant for patient 2 showed completely different macrolide resistance profiles, most striking at position 2058 (Figure 3b). These positions were not among the SNVs clustered into subpopulation structure clusters in patient 2, but *rrl* 2059 was part of cluster 1.D in patient 1, demonstrating that these resistance alleles can arise spontaneously but also persist linked to genetic background.

**Figure 3.**
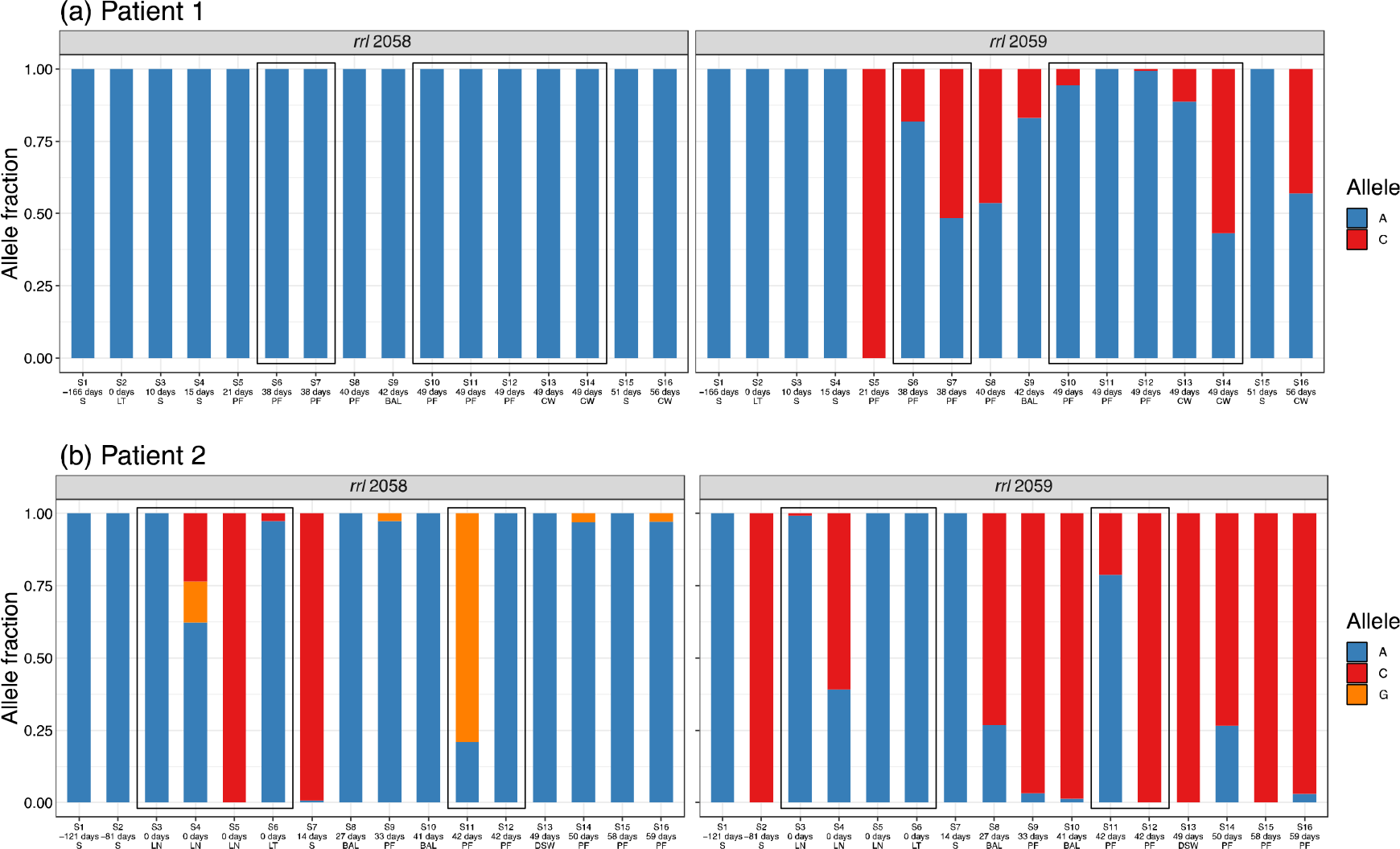
Variants in the *rrl* gene (23S rRNA) arise during treatment and are present in isolates from concurrent samples. Relative allele fractions at these positions show that although the initial sputum isolate was susceptible for both patients, resistance appeared to develop during treatment. Samples are ordered by time, with boxes indicating samples on the same day. Samples taken on the day of transplant are shown in bold text. N.B. Here following the usual convention for the *rrl* gene we use *E. coli* numbering. Positions 2058 and 2059 in *E. coli* correspond to 2269 and 2270 in *M. abscessus.* (S = sputum, BAL = bronchalveolar lavage, CW = chest wound, DSW = drain site swab, LT = lung tissue, LN = lymph node, PF = pleural fluid).

We also observed variable positions in the *rrs* gene in both patients (Figure 4). The *rrs* gene codes for 16S rRNA and is often a site of the emergence of aminoglycoside resistance (e.g. to amikacin), particularly the last few 100bp of the gene where the secondary structure of the rRNA can be affected by multiple mutations^16^. The *de novo* reference assembly for the gene in both patients was identical to the previously characterised sequence from amikacin-resistant *M. abscessus*^16,17^, which was consistent with the measured AMR phenotype of the first sputum sample in Patient 2 but not Patient 1 (Table 3). Subsequently, the patient_1_S2 isolate from lung tissue on the day of transplant had a different allele at position 1174 (C→T; Figure 4), and was partially resistant when phenotyped (Table 3), suggesting this mutation may have been involved in this resistance. In Patient 2, we observed both A and G at position 1374, corresponding to the A1400G mutation which confers high-level amikacin resistance in *M. tuberculosis*^18^. The G allele was dominant by the end of treatment, suggesting that it may have conferred higher resistance and/or been an important compensatory mutation.

**Figure 4.**
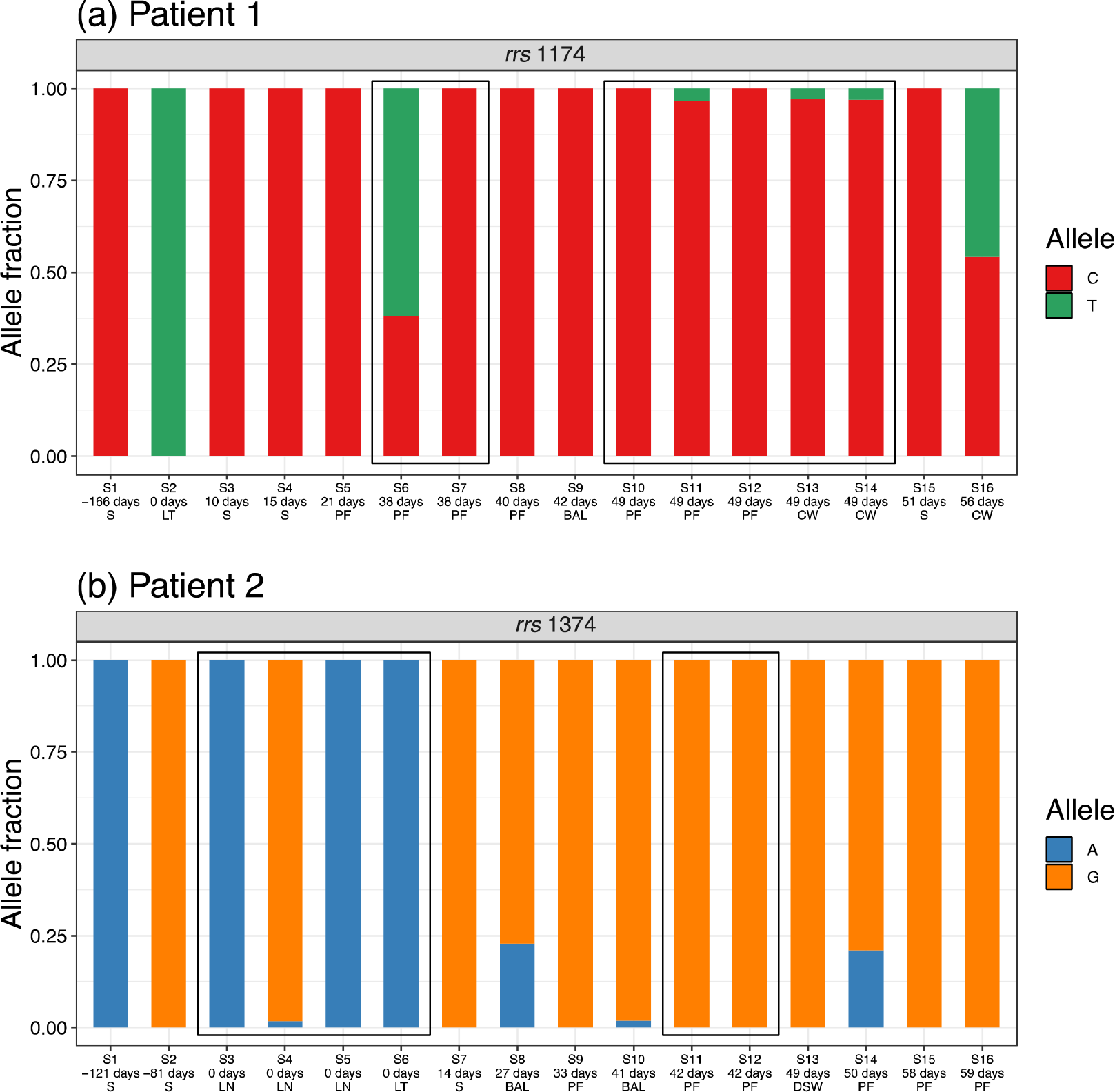
Variants in the *rrs* (16S rRNA) gene over the course of treatment. Patient 1: position 1174. Patient 2: position 1374, previously associated with amikacin resistance (see text). Numbering relative to ATCC 19977 reference. (S = sputum, BAL = bronchalveolar lavage, CW = chest wound, DSW = drain site swab, LT = lung tissue, LN = lymph node, PF = pleural fluid).

Other sites also showed high levels of variation in both patients. After sorting SNVs by the standard deviation of the reference allele fraction across isolates (Supplementary Dataset 1), the most highly variable position in patient 1 was in the putative ferric uptake regulator FurB (MAB_1678c). This variant was present at high abundance in samples 49 days after transplant, with the reference allele fraction only present at <2% in one pleural fluid sample (Supplementary Dataset 1). Ferric uptake regulation has been associated with virulence of pathogenic mycobacteria; in mycobacterial infections the host response deprives the bacteria of iron to prevent replication^19^. Iron is important for growth and virulence in *M. abscessus*^20^ with gallium used as a treatment because of its ability to inhibit iron-dependent enzymes^21^. The most variable position in patient 2 was within a putative linoleoyl-CoA desaturase (MAB_2148), and the second most highly variable position in patient 2 was within the cell division control protein 48 CDC48 (MAB_0347). Population heterogeneity via asymmetric cell division has been suggested as a factor facilitating the survival of *M. tuberculosis* across host physiological niches^22^, and the control of cell division is probably similarly important in the survival of *M. abscessus* across body sites. Both patients had a variable position within *erm(41)* (MAB_2297) although at different positions, which confers inducible resistance to macrolides^23^.

### Sputum samples do not reflect overall within-patient diversity

The first sample from both patients was from sputum. Frequencies of the reference alleles were significantly associated with body site in Patient 2 (Kruskal-Wallis rank-sum test, *p*<0.001) but not Patient 1 (Kruskal-Wallis rank-sum test, *p*=0.13). Reference allele frequencies were significantly higher in subsequent sputum isolates compared to non-sputum isolates for the majority of SNVs in both patients (Patient 1: 51/54 SNVs with *p*<0.05 after Benjamini-Hochberg correction, Patient 2: 61/64 SNVs with *p*<0.05 after Benjamini-Hochberg correction), suggesting that sputum isolates tended to be more similar to the initial sputum isolate used as a reference, even for Patient 1 where 3/3 subsequent sputum isolates were post-transplant (immunosuppressed). This also suggests that non-sputum isolates harboured additional diversity that was not well-sampled using sputum.

## Discussion

In this retrospective study we sought to establish the extent of within-patient variability of *M. abscessus* in two patients who developed severe complications following lung transplant as part of treatment for CF. We used WGS to characterise this variability in isolates from longitudinal clinical samples, and were able to reveal patterns of linkage of SNVs consistent with the presence of multiple subpopulations within patients.

While isolates from the same patient were similar — for example, all inter-isolate distances for the same patient were within the threshold of 25 SNVs previously suggested for inferring potential transmission events^24^ — this does not mean that the variation is not clinically significant. We have demonstrated that within-patient variation can contain biologically and clinically important variation. Notably, we observed that variation at the *rrl* 2058/2059 positions, associated with macrolide resistance, developed over the course of treatment. We observed extensive variation in lung tissue isolates taken from Patient 41 prior to and on the day of transplant (i.e. before the patient was immunosuppressed). Isolates from Patient 1 samples taken prior to and on the day of transplant displayed no variation at these positions, so presumably the phenotypic macrolide resistance reported in these samples at this time was due entirely to inducible resistance conferred by *erm(41)*. Nonetheless, variation at *rrl* 2058/2059 positions was seen in later samples from this patient, suggesting that macrolide use still drives development of high-level macrolide resistance even in the presence of a functional *erm(41)* gene. We also observed variation in the *rrs* gene, another source of resistance to antibiotics targeting ribosomal function (e.g. amikacin), at both previously recorded and novel positions, including an allele that rose in dominance over the course of treatment in Patient 2 (Figure 4).

Based on this data we would question how useful phenotypic testing of isolates recovered from a limited number of sputum isolates is for guiding antimicrobial therapy, as this strategy is unlikely to capture the diversity present in a single sample. Similarly, even though WGS is clearly a valuable tool, if restricted to the analysis of sputum isolates it may also fail to capture an accurate AMR profile of an infection. Previous studies on *M. tuberculosis* have shown that mycobacterial culture reduces the diversity recovered from sputum samples whether the culture was enriched in solid or liquid media^25,26^. It is therefore possible that the diversity found across different sample sites in this study may have been present in the sputum sample, but was possibly lost in the culture step. For *M. tuberculosis*, direct sequencing from sputum samples using capture-based enrichment methods has been shown to recover sample diversity not present in liquid culture^27^. Extending this to this case, it is possible that WGS at high depth applied directly to sputum samples could identify the variants detected between sample sites.

Other highly variable positions in genes involved in e.g. the regulation of ferric uptake (MAB_1678c) and the control of cell division (MAB_0347), are both of direct relevance for the survival of *M. abscessus* across different physiological niches. In a chronic infection, mycobacteria must cope with considerable host stresses, including Fe starvation, which has been highlighted as enabling the persistence of *M. tuberculosis* in granulomas^28^. There is also, of course, phenotypic diversity due to expression. For example, colony morphotype has previously been suggested as providing a distinction between a ‘non-invasive, biofilm-forming, persistent phenotype’ (smooth) and an ‘invasive phenotype that is unable to form biofilms’ (rough), although recent work suggests both morphotypes are capable of aggregation and intracellular survival^29^. Further diversity may come from the subpopulations harboured at different locations within the patient’s lung, which may therefore be an important source of diversity for clinically significant phenotypes.

There is also the possibility that this within-patient variability could lead to different transmission inferences, if different lineages of *M. abscessus* can be present in the lung together. However, the variation found within each of the two patients in this study, who were both infected with a clonal VNTR I strain, did not exceed the variation found between them.

Our findings suggest that the wider diversity present within patients chronically infected with *M. abscessus* is not well-sampled with sputum, and that body site plays a role in subpopulation structure. More widespread sampling of multiple body sites would provide a more accurate picture of the AMR profile of *M. abscessus* infecting a patient, and may be necessary to guide targeted antimicrobial therapy prior to transplant. However, in practical terms this would mean taking biopsies of lung and other tissue, which carries a significant clinical risk and would probably be unfeasible in the patient awaiting transplant. An alternative solution to improve patient management prior to transplant might rely on deep-sequencing of multiple sputum samples. Such a strategy might capture a sufficient fraction of the diversity seen with multiple body site sampling to provide accurate information about the presence of minor variants present in *M. abscessus* infections of CF patients, in particular those conferring AMR.

## Acknowledgements

This project has received funding from the EMPIR programme co-financed by the Participating States and from the European Union’s Horizon 2020 research and innovation programme. The study was also supported by the National Institute for Health Research Biomedical Research Centre at Great Ormond Street Hospital for Children National Health Service Foundation Trust and University College London. LPS and FB acknowledge financial support from the Medical Research Council (grant MR/P007597/1).

## Supplementary data

### Supplementary Dataset 1. SNVs used for clustering in both patients

A list of the variant positions used for clustering to identify subpopulations, sorted in decreasing order of standard deviation across samples. In addition to the position of the variant in the *de novo* assembly reference, its position (if present) in the ATCC 19977 is given, along with the relevant UniProt ID. Samples are sorted in time order.

